# Reef development and Sea level changes drive *Acanthaster* Population Expansion in the Indo-Pacific region

**DOI:** 10.1101/2020.11.18.388207

**Authors:** P.C. Pretorius, T.B. Hoareau

## Abstract

Molecular clock calibration is central in population genetics as it provides an accurate inference of demographic history, whereby helping with the identification of driving factors of population changes in an ecosystem. This is particularly important for coral reef species that are seriously threatened globally and in need of conservation. Biogeographic events and fossils are the main source of calibration, but these are known to overestimate timing and parameters at population level, which leads to a disconnection between environmental changes and inferred reconstructions. Here, we propose the Last Glacial Maximum (LGM) calibration that is based on the assumptions that reef species went through a bottleneck during the LGM, which was followed by an early yet marginal increase in population size. We validated the LGM calibration using simulations and genetic inferences based on Extended Bayesian Skyline Plots. Applying it to mitochondrial sequence data of crown-of-thorns starfish *Acanthaster spp.*, we obtained mutation rates that were higher than phylogenetically based calibrations and varied among populations. The timing of the greatest increase in population size differed slightly among populations, but all started between 10 and 20 kya. Using a curve-fitting method, we showed that *Acanthaster* populations were more influenced by sea-level changes in the Indian Ocean and by reef development in the Pacific Ocean. Our results illustrate that the LGM calibration is robust and can probably provide accurate demographic inferences in many reef species. Application of this calibration has the potential to help identify population drivers that are central for the conservation and management of these threatened ecosystems.

## Introduction

Coral reefs represent one of the most diverse ecosystems on the planet (Hartmann 2018) providing shelter, food, breeding, and spawning for a large diversity of species (Seitz et al. 2014). Coral reefs are also an important resource in terms of fisheries, tourism, and protection from natural disasters (Kunkel et al. 2006; Hoegh-Guldberg et al. 2007). The whole coral ecosystem, therefore, need to be conserved to prevent loss of biodiversity, ecosystem services and utility for dependent coastal communities. The threats associated with global change (e.g. ocean acidification, loss of habitats) have potential impacts on the coral reefs themselves but also on the coastal communities depending on them. Coral bleaching, ocean acidification and overexploitation can all eventually lead to coral death, (Hoegh-Guldberg et al. 2007; Hartmann 2018). In addition, corals may not be able to keep up with rapid increases in sea-level that are predicted to happen (Perry et al. 2018), and which may be worsened by mechanical damages by cyclones (Harmelin-Vivien 1994). This combined effect of all these factors can lead to loss of biodiversity and, a dramatic decrease in fishery stocks (Jones et al. 2004). Moreover, weakened corals are unable to recover from natural disasters like cyclones and are more prone to attack from predators and parasites (Normile 2009; Booth 2011).

The COTS in the genus *Acanthaster* is an obligate corallivore which poses a major threat to corals with outbreaks often leading to large scale devastation of reefs (Baird et al. 2013; Pratchett et al. 2014). Over the last thirty years, many studies have helped advance understanding of the ecology, biology and the genetics of these species, including the drivers of outbreaks (Pratchett et al. 2017). Phylogeographic studies have helped discribe the geographic distribution of genetic diversity and raised four species from the *Acanthaster planci* complex found in the Indo-Pacific region (Vogler et al. 2012; Vogler et al. 2013; Haszprunar et al. 2017). These studies provide multiple large genetic datasets, which are complemented by ecological data that help a better understanding of the factors that may affect larval and adult behaviour as well as larval development. This means that the COTS is a useful study model, which may aid in understanding how reef dependent species responded to post-LGM change in marine environments. Studying this species may help with future predictions and management of these ecosystems.

Many genetic methods can be used to infer the demographic history of species. These methods can be broadly divided into mismatch analyses, neutrality tests, maximum likelihood and Bayesian methods. Each of these methods has unique limitations regarding the information that can be gleaned from the number and type of parameters considered, from limitations enforced by data availability, and from calculation efficiency and confounding factors (Felsenstein 1992; Ramos-Onsins and Rozas 2002; Beerli 2006; Ho and Shapiro 2011; Li and Durbin 2011). Among these methods, the Bayesian Skyline Plot has been extensively used for the reconstruction of effective population size through time especially using mitochondrial DNA data (Heled and Drummond 2008; Ho and Shapiro 2011). Although some studies highlighted potential issues when applied on datasets from admixed populations (Heller et al. 2013), it remains otherwise a robust method that can help testing hypotheses on climatic factors that could have impacted the demographic history of targeted species (Ho and Shapiro 2011).

In retracing population histories, the resolution of genetic inferences depends on the amount of data used but the accuracy of the timeline relies on the correct calibration of the molecular clock. Methods of calibration most often depend on fossils and biogeographic events and sometimes use radiocarbon datings of ancient DNA. Fossil calibration is the most prevalent of these methods (Hipsley and Müller 2014; Ho et al. 2015) and in conjunction with biogeographic dating (e.g. Lessios 2008) rely on events that are most often at least 1 million years old. This results in slow substitution rates that overestimate the time and population parameters at intraspecific levels (Ho et al. 2011; Crandall et al. 2012). The use of ancient DNA provides an ideal alternative (Orlando and Cooper 2014). It involves using sequences from samples that are independently dated with radiocarbon or stratigraphic methods to date the tips of phylogenetic trees (Shapiro et al. 2011). The use of ancient DNA is however limited by the availability of well-preserved tissues and its utility is limited for marine taxa, especially for tropical species (Crandall et al. 2012).

The calibration based on expansion dating (Crandall et al. 2012) improves upon previous methods by using two calibration points of possible expansion at 19.6 kya and 14.6 kya using ecological assumptions. This method was specifically developed for species occurring on the Sunda Shelf and is constrained by the Two-Epoch-Model which makes it difficult to generalise to other reef species. The CDT method (Hoareau 2016) improves on the expansion dating method by using the full variance of both environmental and population transition parameters to estimate a more accurate mutation rate, which provides more accurate demographic timelines. The CDT is however developed with ecological assumptions of temperate species (Hoareau 2016), which may make it unsuitable for reef species.

To accurately retrace the demography of reef species, we developed the LGM calibration method, a new method that accounts for reef species paleoecology since the LGM. We validated the calibration method using a simulation study, and then applied it to different populations of the iconic *Acanthaster* species in the Indo-West Pacific region. We then designed a similarity index to better evaluate the respective role of sea-level changes and modern coral development on their demography.

## Material and methods

### Simulated and empirical datasets

To validate the calibration methods we simulated different demographic scenarios using FASTSIMCOAL v2 (Excoffier et al. 2013). The scenarios are characterised by multiple changes in population size that mimic actual environmental variations like changes in Relative Sea Level (Waelbroeck et al. 2002). We simulated a demographic scenario using a transformed RSL as a proxy (table S1) with 100 sets of 100 sequences of 2500 bp in length setting the effective population size to ten million (10×10^6^) with a single deme and no migration.

We recovered sequence datasets of COTS from the Indian and Pacific Oceans (Vogler et al. 2012; Vogler et al. 2013) that comprised mitochondrial DNA (mtDNA) sequences of the cytochrome oxidase subunit 1 (*COI)* and the hypervariable region (D-loop) of the control region (*CR*). To avoid potential biases that could result from genetic structure (Heller et al. 2013), we used samples representing homogenous regional populations with at least 20 individuals as identified by (Vogler et al. 2012; Vogler et al. 2013). These populations that were originally part of the *A. planci* complex have recently been suggested to belong to distinct species (Haszprunar et al. 2017): *A. mauritiensis* (Southwestern Indian Ocean, N=79), *A. planci* (north-eastern Indian Ocean, N=54), *A. planci* (north-western Indian Ocean, N=24), *A. solaris* (western Pacific, N=584), *A. solaris* (Central Pacific, N=66) (Table 1; Fig. 1). To avoid including individuals that violate the assumption of panmixia we exclusively used regional populations that were identified to be significantly different from others in each major area (NIO, SIO, WP, CP, EP) taking care to exclude haplotypes found in more than one region.

**Table 1.**
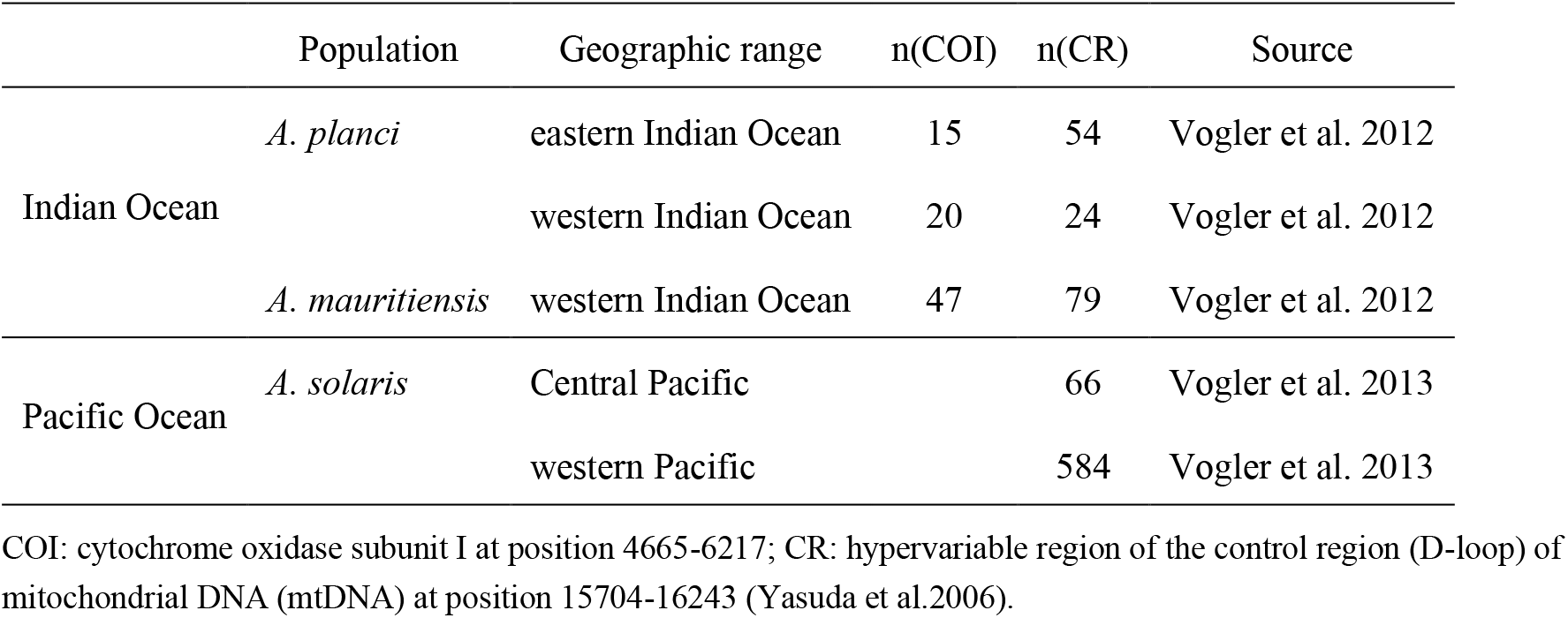
Details of the sequence datasets of the five populations considered in the genus *Acanthaster*.

|  | Population | Geographic range | n(COI) | n(CR) | Source |
| --- | --- | --- | --- | --- | --- |
| Indian Ocean | *A. planci* | eastern Indian Ocean | 15 | 54 | Vogler et al. 2012 |
|  |  | western Indian Ocean | 20 | 24 | Vogler et al. 2012 |
|  | *A. mauritiensis* | western Indian Ocean | 47 | 79 | Vogler et al. 2012 |
| Pacific Ocean | *A. solaris* | Central Pacific |  | 66 | Vogler et al. 2013 |
|  |  | western Pacific |  | 584 | Vogler et al. 2013 |

COI: cytochrome oxidase subunit I at position 4665-6217; CR: hypervariable region of the control region (D-loop) of mitochondrial DNA (mtDNA) at position 15704-16243 (Yasuda et al.2006).

**Fig. 1.**
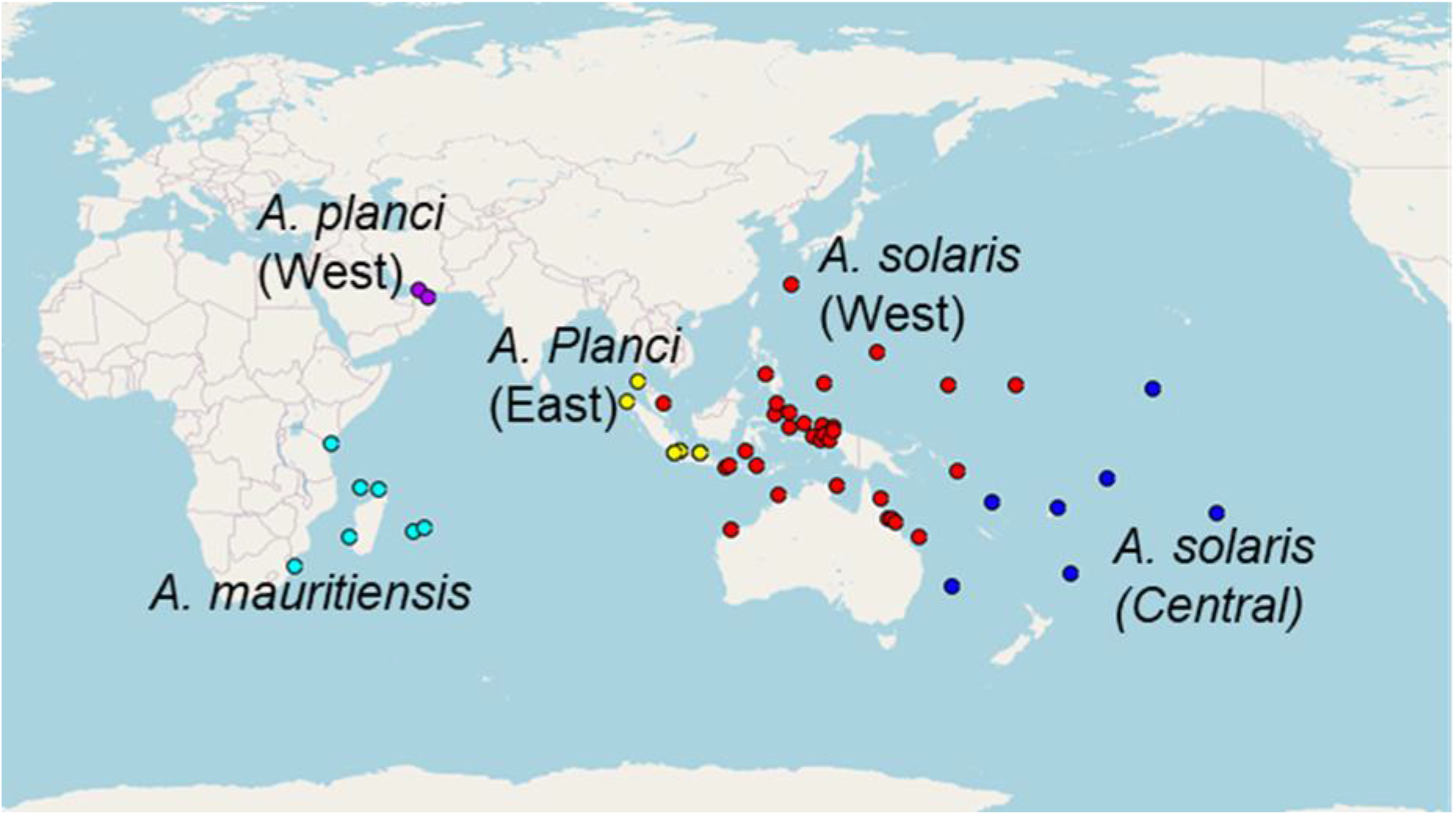
Sampling locations of the five *Acanthaster spp* in the Indo-Pacific region

### Genetic diversity and test of demographic expansion

When available for each individual, *COI* and *CR* were concatenated for downstream analyses. We aligned the sequences using the online version of MAFFT v7.0 (Katoh et al. 2002), applying the L-INS-i iterative refinement method, with a gap penalty set to 1.53, the BLOSUM62 scoring matrix and the default nucleotide scoring matrix of 200PAM. We calculated standard summary statistics for each population using DNASP v5.1 (Librado and Rozas 2009). These included nucleotide diversity (*π*), haplotype diversity (*Hd*), and three population parameters that help identify demographic changes, which include Tajima’s *D*, Fu’s *F*_*S,*_ and Ramos-Onsins and Rozas’ *R*_*2*_ (Ramos-Onsins and Rozas 2002). Population expansion is indicated by significantly negative values for *D* and *F*_*S*_, and significantly small values for *R*_*2*_ (Ramos-Onsins and Rozas 2002). We also generated a mismatch distribution for each population using DNASP v5.1 and compared observed and expected distributions for both cases of population expansion and constant size. We supplemented these comparisons by testing for deviation from the constant population size curve generated by DNASP v5.1 using a *χ*^2^ test.

### Inference of population history using coalescent-based approaches

We reconstructed the demographic history of each population using the extended Bayesian skyline plots model (EBSP) (Heled and Drummond 2008) implemented in BEAST v2.6.2 (Bouckaert et al. 2019). We selected the gamma site model with the HKY nucleotide substitution model and empirical frequency estimation. In the first analyses, we kept the clock rate value as 1.0 to express estimates of population and time parameters in terms of mutations per site. We set a strict clock model and changed the mean population size hyper-prior from *1/x* to a normal distribution, as suggested by the developers to improve the rate of convergence (Heled and Drummond 2008; Bouckaert et al. 2014). For each run, we ran 100 million Markov Chain Monte Carlo iterations while logging every 10,000 to obtain a total of 10,000 samples of genealogies, and population and time parameters. We used TRACER v1.7.1 (Rambaut et al. 2018) to verify that the sampling scheme of the runs was correct based on a threshold for the effective sample sizes (ESSs) superior to 200. After discarding the first 10% of the samples as burn-in, we generated a csv output file for the skyline plots using EBSPANALYSER v2.5.1 (beast2.org/ebspanalyser/). These analyses were repeated at a later stage applying the different mutation rates calculated independently (see the section on molecular clock calibration).

### Calibration of the molecular clock

To transform evolutionary times into calendar years, we used three different calibration methods: biogeographically dated divergence (Lessios 2008), Calibration by Demographic Transition (CDT; Hoareau 2016), and a new calibration specifically designed for tropical coastal species. The first calibration method uses the closure of the Isthmus of Panama to calibrate the divergence of assumed sister species (Lessios 2008). We used the average mutation rate obtained for mitochondrial protein-coding genes in echinoderms (Lessios 2008). The CDT method was developed to account for the association between environmental factors and the demography of wild populations (Hoareau 2016), and we applied this method for additional comparison.

We also developed a new calibration method that assumes that coral reef species experienced a population bottleneck during the LGM and post-LGM expansion. This assumption is likely as a very large fraction of tropical species show signs of genetic bottleneck or local extinction during the LGM (26.5-19 kya; McCabe et al. 2009) (Paulay 1990; Grant 2015; Ludt and Rocha 2015) and the improvement of environmental conditions at the end of the LGM likely led to an expansion in reef populations (Ludt and Rocha 2015). Based on these assumptions, we consider that the first 5% increase in population size should coincide with the end of the LGM and the first 5% increase in RSL (17.775 kya). We call this method as the Last Glacial Maximum Calibration.

The time and population parameters (mean, median, and 95% confidence interval values) obtained from the EBSP outputs were calibrated using the three mutation rates and compared (Fig. S2). This provided time estimates in calendar years and estimates of female effective population size.

### Validation and cross-checking of the new calibration method using environmental factors

We used simulated populations to validate the LGM calibration by visual comparison of the inferred histories after calibration to a simulated scenario. We then compared the simulated RSL scenario in conjunction with a reef development curve (Hoareau and Pretorius in prep) to the newly inferred demographic histories of the populations of *Acanthaster* species using a newly developed Similarity Index which was inspired by the recent Dissimilarity Index used to compare BSP curves (Miller and Amos 2018). To account for the differences in the scale of expansion, we standardized the range of *Ne τ* from 0 to 1 which allows the comparison of the distances between curves across a selection of points. We calculated the distance between the relative median of the EBSP plots and the demographic proxy of the simulation and a reef development curve at the closest corresponding time estimates based on the simulated scenario. The relative contribution of RSL and reef development was altered in 25% intervals starting at 0% for one and 100% for the other which provided a gradient of possible scenarios (Fig. S4) to which we calculated the distances previously mentioned. The sum of all these distances yield the SI of that population-scenario comparison, which is calculated as follows:

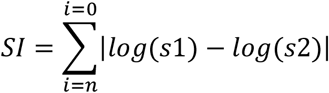

Where i is the number of time estimates in the demographic inferences based on simulated data, s1 the relative population size at that point in time and s2 the relative population size estimate at the closest match point in time. This allows a relative comparison of demographic histories to driving factor scenarios based on the timing of expansions and shape of the inferred EBSP. In addition, we used the standardized curves to estimate growth rates (Hoareau 2020) with:

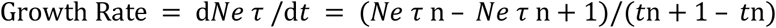

Which may help to identify inflexion points in the demographic histories. In addition, we assessed the impact of the calibration used on the Similarity of the CDT simulations (Hoareau 2016) and assessed the overall similarity and variance.

## Results

The application of the LGM calibration method to 100 simulated sequence dataset provided calibrated genetic inferences that closely followed the RSL curve initially used as a demographic scenario. We obtained the expected patterns of expansion after the LGM even though there is a wide variation in the TMRCA values (20-69 kya) and the level of expansion (Fig. 2).

**Fig. 2.**
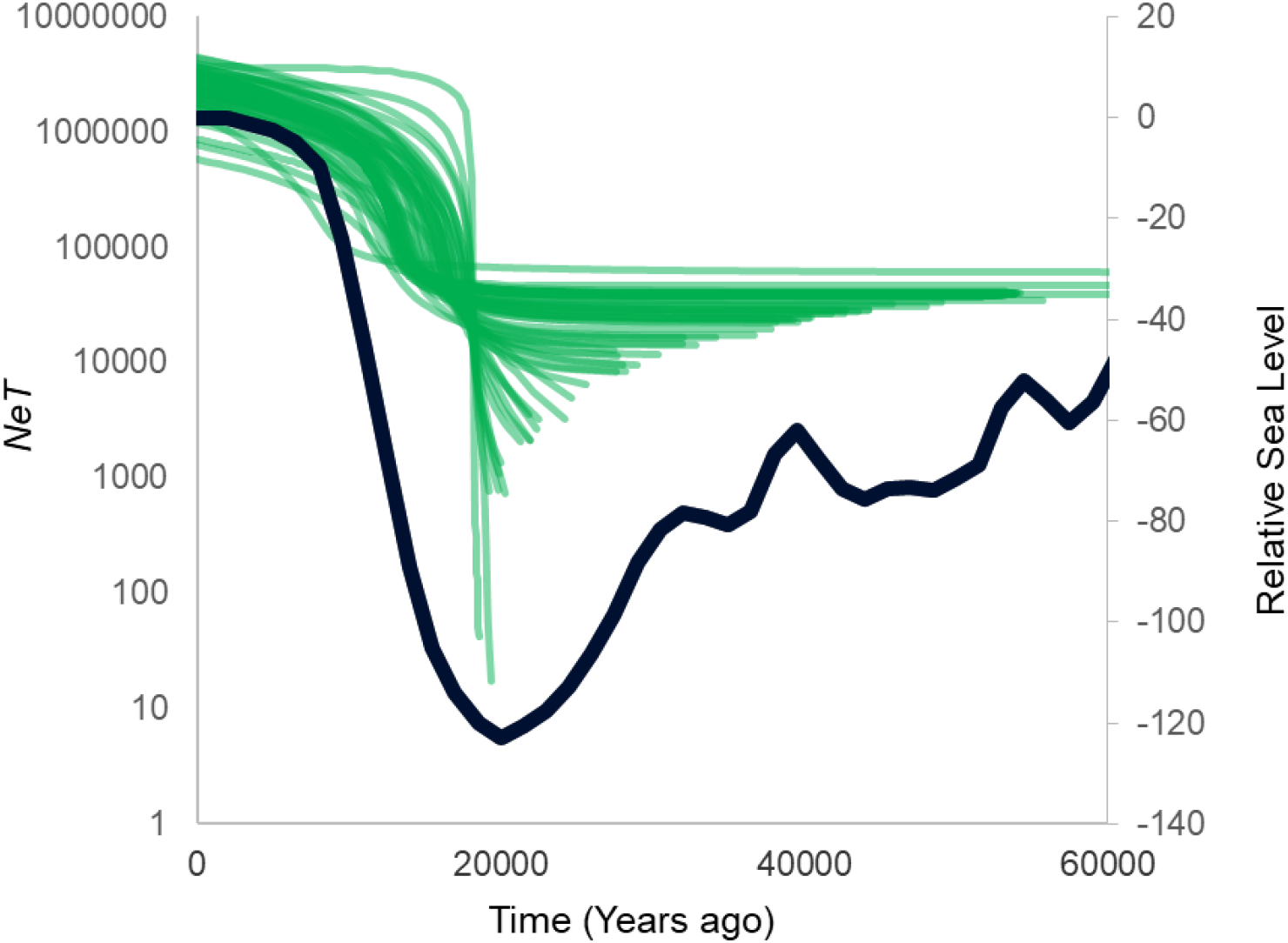
Inferences based on the Extended Bayesian Skyline Plots model illustrated by the median *Ne* values of 100 inferred demographic histories (green curves). The inferences are performed on sequence datasets based on simulations and are calibrated by use of the new LGM calibration. The dark curve represents the Relative Sea Level (Waelbroeck et al. 2002) and has been used here as a demographic proxy for reef species.

A pattern of expansion is visible in the EBSP plots for all populations analysed (Table 2, Fig. 3, Fig. S2). In addition, Likelihood-based methods derived from the EBSP results show deviation from a constant population size model for all populations except *A. solaris* from the Central Pacific (Table 2). Mismatch distributions show a unimodal distribution that differs significantly from constant population size distributions which suggest population expansion in all populations except *A. solaris* population from the Central Pacific (bimodal distribution; Fig. S3). Tests based on statistical parameters (*π*, *Hd*, *D*, *FS* and *R2*) only show significant expansion for *A. solaris* from the western Pacific and *A. mauritiensis* (Table 2).

**Table 2.** Summary statistics and Neutrality tests based on the mitochondrial control region and cytochrome oxidase subunit 1 sequence data for the identified populations of *Acanthaster.* Significant values are highlighted (*).

| Marker | Population | *N* | *H* | *Hd* | $\pi$ | *D* | *F<sub>s</sub>* | *R<sub>2</sub>* |
| --- | --- | --- | --- | --- | --- | --- | --- | --- |
| CR | *A. solaris* WP | 584 | 295 | 0.99 | 0.02 | -2.07* | -565.79* | 0.02 |
|  | *A. solaris* CP | 66 | 50 | 0.99 | 0.04 | -0.40 | -19.04 | 0.09 |
|  | *A. mauritiensis* SWIO | 79 | 57 | 0.98 | 0.02 | -1.62 | -49.14* | 0.05 |
|  | *A. planci* NWIO | 24 | 12 | 0.91 | 0.004 | -1.06 | -6.55 | 0.09 |
|  | *A. planci* NEIO | 95 | 35 | 0.97 | 0.01 | -1.06 | -24.67 | 0.07 |
| COI | *A. mauritiensis* SWIO | 51 | 13 | 0.53 | 0.002 | -2.17* | -11.07* | 0.04 |
|  | *A. planci* NWIO | 19 | 3 | 0.21 | 0.0003 | -1.51 | -1.80 | 0.15 |
|  | *A. planci* NEIO | 16 | 3 | 0.24 | 0.0006 | -1.70 | -0.90 | -0.18 |

COI: cytochrome oxidase subunit I at position 4665-6217; CR: hypervariable region of the control region (D-loop) of mitochondrial DNA (mtDNA) at position 15704-16243 (Yasuda et al.2006).

**Fig. 3.**
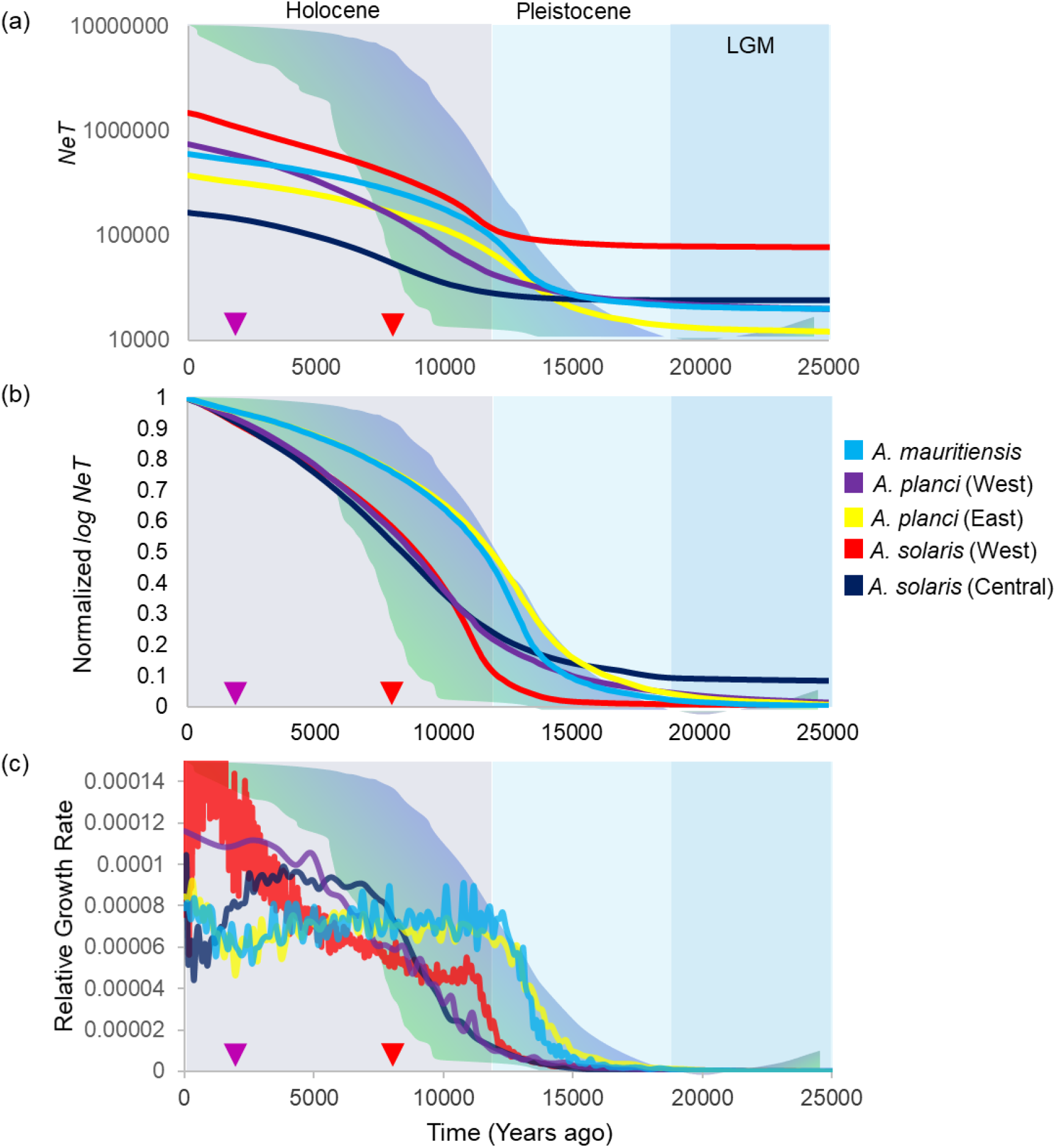
Demographic reconstructions in five *Acanthaster* populations applying the LGM calibration and major environmental factors that are assumed to have impacted reef species. **(a)** Median values of *NeT* for the five *Acanthaster* populations in the Indo-Pacific region overlaid with relative sea-level change (blue) and reef development (green). **(b)** Inferred demographic history of 5 *Acanthaster* populations in the Indo-Pacific region overlaid on the gradient of possible scenarios with changing relative contributions from reef development (green) and sea-level change (blue). **(c)** Relative growth rates of the different *Acanthaster* populations compared to the gradient of scenarios. The red arrow indicates the oldest *A. solaris* ossicles on the eastern Great Barrier Reef (8 kya) and the pink arrow indicates the start of dramatic increases in *A. solaris* ossicle density in the same region (2 kya).

The mutation rates we obtained using the LGM calibration vary between the five *Acanthaster* populations but are very high, ranging from 3.67×10^−8^ to 8.90×10^−7^ (mutations per site per year; Table 3). The two populations of *A. solaris* from the Indo-Pacific have much faster rates, but these estimates are only based on *CR* sequences unlike populations from the Indian Ocean that combine both *CR* and *COI* sequences. Rates obtained by CDT calibration are generally slower than LGM calibration rates, but the trend of faster rates in *A. solaris* populations remains (Table 3).

**Table 3.** Calibration rates obtained for LGM and CDT calibration methods on the five populations of *Acanthaster* species.

| Population | LGM | CDT |
| --- | --- | --- |
| *A. planci* eastern Indian Ocean | $1.46 \times 10^{-07}$ | $1.20 \times 10^{-07}$ |
| *A. planci* western Indian Ocean | $4.75 \times 10^{-08}$ | $7.69 \times 10^{-08}$ |
| *A. mauritiensis* | $3.67 \times 10^{-08}$ | $1.19 \times 10^{-07}$ |
| *A. solaris* Central Pacific | $8.90 \times 10^{-07}$ | $6.74 \times 10^{-07}$ |
| *A. solaris* western Pacific | $4.29 \times 10^{-07}$ | $3.18 \times 10^{-07}$ |

Rates are expressed in mutations per site per year

The growth rate of the populations varies greatly, with *A. solaris* from western and central Pacific showing 15- and 7-fold expansions, respectively. The populations of *A. planci* from northwestern- and eastern Indian Ocean show 31- and 32-fold expansions, respectively, whilst *A. mauritiensis* experienced a 40-fold expansion (Fig. 3). The timing of expansion also varies among populations and even among those belonging to the same species (Fig. 3). Populations of both *A. solaris* and *A. planci* from the north-western Indian Ocean show more recent expansions compared to *A. mauritiensis* and *A. planci* from the north-eastern Indian Ocean (Fig. 3B). These observations are clearer on the growth patterns, which peak earlier for *A. mauritiensis* and *A. planci* (north-eastern Indian Ocean) compared to the other populations (Fig. 3C). The western Pacific population of *A. solaris* shows two distinct phases of exponential growth, which contrasts with the single increase observed in other populations. Spikes of growth rate in this population roughly correspond to an increase in abundance of COTS ossicle on the Great Barrier Reef (Fig. 3; Walbran et al. 1989).

Analyses based on the specifically developed similarity index confirm that each population have unique demographic histories. These analyses indicate that the two EBSP curves for the populations of *A. solaris* matched modern reef development during the Holocene whilst EBSP curves of *A. mauritiensis* and *A. planci* (north-eastern Indian Ocean) were more similar to sea-level changes that occur since the LGM. The EBSP curve of *A. planci* population from the north-western region of the Indian Ocean was equally similar to RSL and Reef development curves.

## Discussion

### Validation of the LGM calibration method

The LGM calibration method developed in the present work assumes that reef species have experienced a bottleneck during the LGM and have had an increase in population size during the deglaciation (Hoareau and Pretorius in prep.). These assumptions should hold for most reef species considering that the carrying capacity of reef environments (available resource and coastal habitats) were at a minimum during the LGM (Ludt and Rocha 2015). These harsh conditions led to local extinctions of some species (Paulay 1990), and most likely resulted in strong bottlenecks in species that survived (Ludt and Rocha 2015). When species experience a strong bottleneck, previous genetic signatures are lost, and only post-bottleneck demography remains (Grant and Cheng 2012; Grant 2012; Hoareau and Pretorius in prep.). Following these observations and the assumptions of LGM-bottlenecks in reef species, we considered that the first 5% increase in inferred *Ne* coincides with an equivalent 5% increase in RSL, which occurred at 17.8 kya. This date also coincides with the end of the first post-glacial Meltwater Pulse 1Ao (19.6-18.8 kya), which is the first post-LGM rapid sea level increase (Warrick et al. 2012). We obtained genetic inferences that were all consistent with the original simulated scenario (Fig. 2), suggesting that the LGM calibration is robust. These results together with the assumptions of LGM-bottleneck in reef species suggest that the LGM calibration method is suitable to infer demographic histories for a large range of species depending on reef ecosystems. This should help address the disconnect observed between environmental drivers, hard evidence and genetic inferences for coral reefs highlighted in a recent study (Hoareau and Pretorius in prep).

### LGM calibration rates are faster than rates derived from biogeographic divergence

The mutation rates obtained from the LGM calibration method are high, ranging from 3.67×10^−8^ – 8.9×10^−7^ changes per site per year; Table 3) with *A. solaris* populations showing rates that are from 3 to 24-fold faster than other populations. The mutation rates derived from the LGM calibration are considerably faster than substitution rates obtained for echinoderms and derived from divergences resulting from the closure of the Isthmus of Panama (1.9×10^−8^ per site per year; Lessios et al. 2008). Most of the LGM calibration rates are of the same order than rates obtained using other demographic calibrations in other marine invertebrates, including 4.6×10^−8^ – 1.34×10^−7^ per site per year for the expansion dating method (Crandall et al. 2012) and 1.35×10^−7^ per site per year for the CDT method (Hoareau 2016). Applying the CDT method to *Acanthaster* populations returns rates that are close to values obtained for the LGM calibration (Table 3; 7.67×10^−8^ – 6.74×10^−7^ per site per year).

The higher perceived rates in *A. solaris* populations may be the result of the use of both *CR* and *COI* markers instead of just the *COI* for the rest of the populations. These markers are different, and *CR* is often considered to be the region with the highest substitution rate in the mitogenome, including in echinoids (Bronstein and Haring 2018). Higher rates in *Acanthaster* compared to other invertebrates and even between species may be tied to differences in ecology and other biological traits like fecundity. Long-lived species with lower fecundity tend to have slower substitution rates and vice versa (Wu and Li 1985; Martin and Palumbi 1993; Baer et al. 2007; Bromham 2011; Hoareau 2016). Based on this assumption, faster rates observed in *Acanthaster* populations could be caused by a very large fecundity, and this is supported by the fact that each female can produce millions of eggs, which partly explain the scales of the outbreaks (Pratchett et al. 2014). The fecundity is believed to be higher in *A. solaris*, which is more prone to outbreaks (Haszprunar et al. 2017; Pratchett et al. 2017) when compared to the other two species, *A. mauritiensis* and *A. planci*. Even if some of these rates are unusually high, application of these rates leads to genetic inferences that are ecologically sound as it matches the assumptions of an LGM bottleneck and late glacial expansion (Fig. S2) previously identified (Ludt and Rocha 2015; Hoareau and Pretorius in prep.). By using a single point calibration rather than the calibrating the whole expansion as in previous studies (Crandall et al 2012; Hoareau 2016), the new method provides individually tailored rates while reducing the constraints on the expansion. The LGM calibration therefore allows for the identification of drivers that affect the demography following the LGM, even during the expansion phase.

### Environmental drivers of *Acanthaster* demography

A range of evidence indicates that all populations of *Acanthaster* have expanded in the past. The EBSP plots indicate that all populations show an increase in size (Fig. S2), which is confirmed by likelihood ratio tests, mismatch distributions and neutrality tests that indicate a deviation from constant size (Fig. S3). The lack of significance in some neutrality tests may be a result of small sample sizes, which has been identified in the past on other echinoderms (Hoareau personal observation). This may explain why results are significant in *A. solaris* (n=584 sequences) and *A. mauritiensis* (n=126 sequences) but not in the other populations (n=44, 66 and 69).

As expected by the LGM calibration and the ecological assumptions for *Acanthaster* (Adkins, McIntyre and Schrag, 2002; Clark et al. 2009; Pratchett et al. 2017; Ludt et al. 2015; Deaker et al. 2020; Hoareau and Pretorius in prep.), the expansion consistently occurs after the LGM and within the same broad timeframe for all populations (Fig. 3, Fig. S2). These expansions also vary in intensity, with increase ranging from 7 to 15-fold for Pacific populations and 30 to 40-fold for populations from the Indian Ocean. Ultimately these calibrations offer more realistic demographic reconstructions of reef species that are in line with ecological factors that are independently known to affect them when compared to what has been observed in the recent past (Hoareau and Pretorius in prep).

To identify whether sea-level fluctuations and coral reef development are potential drivers, we evaluated and compared the curves of RSL (Waelbroeck et al. 2002) and reef development (Hoareau and Pretorius in prep) to the calibrated reconstruction of population size of *Acanthaster* populations. To do so, we used a specifically designed similarity index inspired by the dissimilarity measure from a recent study on human populations (Miller and Amos 2018). Applying this index, the two populations from the Pacific Ocean seem to be more influenced by reef development (Fig. 4), whilst *A. mauritiensis* and the population of *A. planci* from the northeastern Indian Ocean are more directly associated with sea-level changes. The population of *A. planci* from the northwestern Indian Ocean seems to be equally impacted by sea-level changes and reef development.

**Fig. 4.**
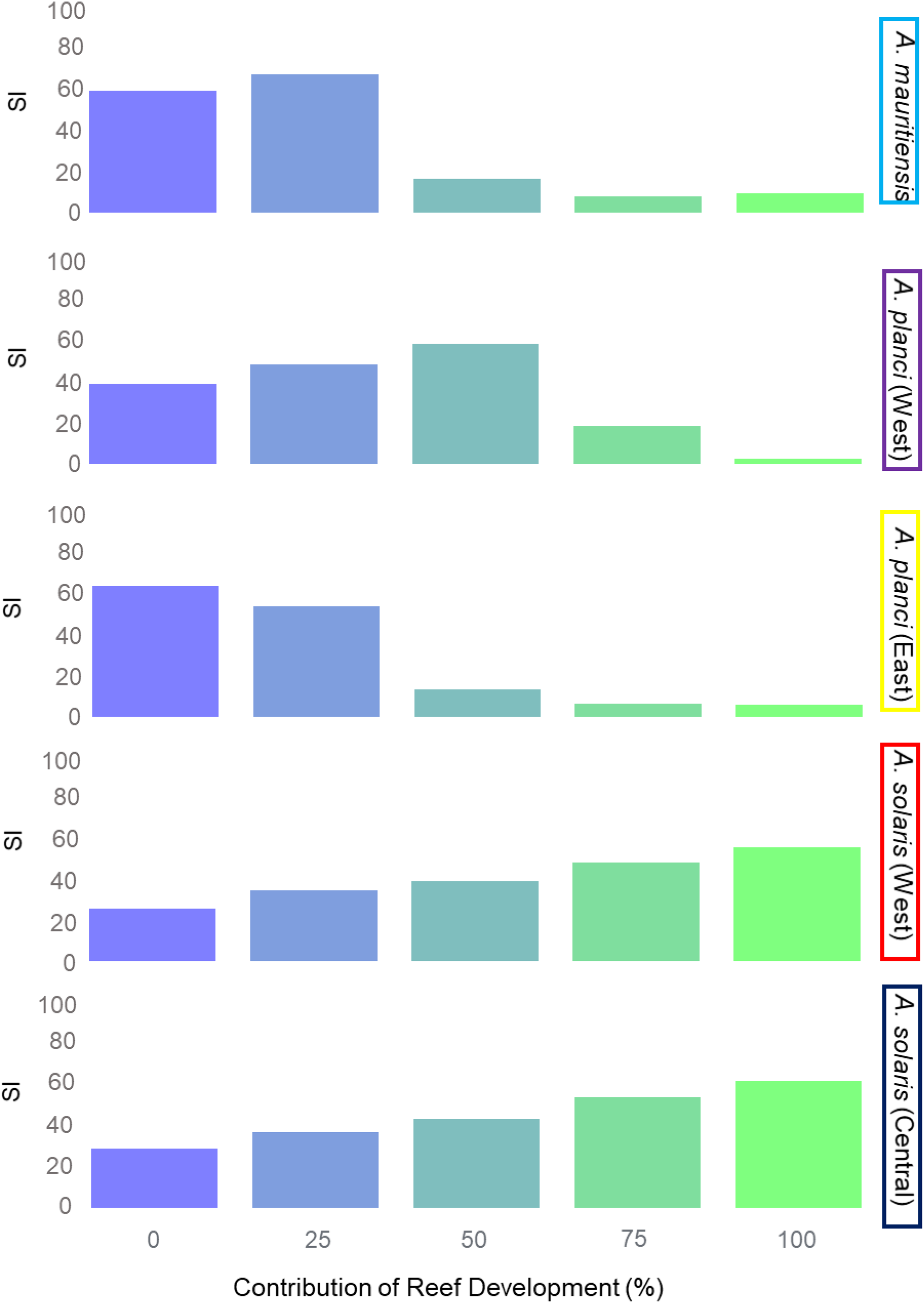
Similarity Index of the inferred demographic histories of each of the five *Acanthaster* species compared to the possible gradient of scenarios. These scenarios are based on reef development and changes in relative sea level. The scenarios are generated by 25% interval combinations of the change in relative sea level and reef development. 0% represents a curve matching the RSL curve, whilst 100% represents a curve matching the coral reef development

Despite being developed for reef species, the present study represents a proof of concept for other study models from more specific environments. It represents a framework to provide appropriate calibrations and inferences that allow for correct identification of demographic drivers, which will eventually help target management and conservation efforts. New questions arise regarding factors that affect the calibration rate, and although some considerations have been given to the dates used and the intensity of the bottleneck (Hoareau and Pretorius in prep), simulations are important tools to further assess the effect of heterogeneity of substitution rates between markers or the impact of using multiple markers on the resolution of inferences.

## Acknowledgements

We thank the Centre for High Performance Computing for providing the resource needed for performing the analyses in this study. TBH was supported by the University of Pretoria’s senior postdoctoral programme.

## References

Adkins JF, McIntyre K, Schrag, DP (2002) The salinity, temperature, and δ18O of the glacial deep ocean. Science 298: 1769–1773.

Baer CF, Miyamoto MM, Denver DR (2007) Mutation rate variation in multicellular eukaryotes: Causes and consequences. Nat. Rev. Genet. 8:619–631

Beerli P (2006) Comparison of Bayesian and maximum-likelihood inference of population genetic parameters. Bioinformatics 22: 341–345

Booth H (2011) Effects of Coral Stressing on the Feeding Preferences of the Coral Predator, Acanthaster Planci. Thesis, Independent Study Project (ISP) Collection

Bouckaert R, Heled J, Kühnert D, Vaughan T, Wu CH, Xie D, Drummond AJ (2014) BEAST 2: a software platform for Bayesian evolutionary analysis. PLoS Comput Biol 10: e1003537

Bromham L (2011) The genome as a life-history character: Why rate of molecular evolution varies between mammal species. Phil. Trans. R. Soc. Lond. B Biol. Sci. 366:2503–2513

Bronstein O, Kroh A, Haring E (2018) Mind the gap! The mitochondrial control region and its power as a phylogenetic marker in echinoids. BMC evolutionary biology 18: 80

Clark PU, Dyke AS, Shakun JD, Carlson AE, Clark J, Wohlfarth B, McCabe AM (2009) The last glacial maximum. Science 325: 710–714

Crandall ED, Sbrocco EJ, DeBoer TS, Barber PH, Carpenter KE (2012) Expansion dating: calibrating molecular clocks in marine species from expansions onto the Sunda Shelf following the Last Glacial Maximum. Molecular Biology and Evolution 29: 707–719

Deaker DJ, Agüera A, Lin HA, Lawson C, Budden C, Dworjanyn SA, Byrne M (2020) The hidden army: corallivorous crown-of-thorns seastars can spend years as herbivorous juveniles. Biology Letters 16: 20190849

Delrieu-Trottin E, Mona S, Maynard J, Neglia, V, Veuille M, Planes S (2017) Population expansions dominate demographic histories of endemic and widespread Pacific reef fishes. Scientific reports 7: 1–13

Drummond AJ, Rambaut A, Shapiro B, Pybus OG (2005) Bayesian coalescent inference of past population dynamics from molecular sequences. Molecular biology and evolution 22: 1185–1192

Excoffier L, Dupanloup I, Huerta-Sánchez E, Sousa VC, and Foll, M (2013) Robust demographic inference from genomic and SNP data. PLoS Genet 9: e1003905

Felsenstein J (1992) Estimating effective population size from samples of sequences: inefficiency of pairwise and segregating sites as compared to phylogenetic estimates. Genetics Research: 59: 139–147

Grant WS (2015) Problems and cautions with sequence mismatch analysis and Bayesian skyline plots to infer historical demography. Journal of Heredity 106: 333–346

Harmelin-Vivien ML (1994) The effects of storms and cyclones on coral reefs: a review. Journal of Coastal Research: 211–231

Hartmann A (2018) Can Baby Corals Improve the Reefs of Tomorrow. Available at: https://hmnh.harvard.edu/baby-corals.

Haszprunar G, Vogler C, Wörheide G (2017) Persistent gaps of knowledge for naming and distinguishing multiple species of crown-of-thorns-seastar in the Acanthaster planci species complex. Diversity 9: 22

Heled J, Drummond AJ (2008). Bayesian inference of population size history from multiple loci. BMC Evolutionary Biology 8: 289

Heller R, Chikhi L, Siegismund HR (2013) The confounding effect of population structure on Bayesian skyline plot inferences of demographic history. PloS one 8: e62992

Hipsley CA, Müller J (2014) Beyond fossil calibrations: realities of molecular clock practices in evolutionary biology. Frontiers in genetics 5: 138

Hoegh-Guldberg O, Mumby PJ, Hooten AJ, Steneck RS, Greenfield P, Gomez E, Knowlton N (2007) Coral reefs under rapid climate change and ocean acidification. Science 318: 1737–1742

Ho SY, Lanfear R, Bromham L, Phillips MJ, Soubrier J, Rodrigo AG, Cooper A (2011) Time-dependent rates of molecular evolution. Molecular ecology 20: 3087–3101

Ho SY, Shapiro B (2011) Skyline-plot methods for estimating demographic history from nucleotide sequences. Molecular ecology resources 11: 423–434

Ho SY, Tong KJ, Foster CS, Ritchie AM, Lo N, Crisp MD (2015) Biogeographic calibrations for the molecular clock. Biology letters 11: 20150194

Hoareau TB (2016) Late glacial demographic expansion motivates a clock overhaul for population genetics. Systematic Biology 65: 449–464

Hoareau TB (2020. Whales and Men: genetic inferences uncover a detailed history of hunting in bowhead whale. bioRxiv [doi: https://doi.org/10.1101/2020.04.09.033191]

Hoareau TB, Pretorius PC (2020) Glacial cycles drive the contraction-expansion dynamics of reef species. Unpublished Manuscript

Jones GP, McCormick MI, Srinivasan M, Eagle JV (2004) Coral decline threatens fish biodiversity in marine reserves. Proceedings of the National Academy of Sciences 101: 8251–8253

Katoh K, Misawa K, Kuma KI, Miyata T (2002) MAFFT: a novel method for rapid multiple sequence alignment based on fast Fourier transform. Nucleic acids research 30: 3059–3066

Kunkel CM, Hallberg RW, Oppenheimer M (2006) Coral reefs reduce tsunami impact in model simulations. Geophysical research letters 33: L23612

Lessios HA (2008) The great American schism: divergence of marine organisms after the rise of the Central American Isthmus. Annual Review of Ecology, Evolution, and Systematics 39: 63–91

Li H, Durbin R (2011) Inference of human population history from individual whole-genome sequences. Nature, 475: 493–496

Librado P, Rozas J (2009) DnaSP v5: a software for comprehensive analysis of DNA polymorphism data. Bioinformatics 25: 1451–1452

Ludt WB, Rocha LA (2015) Shifting seas: The impacts of Pleistocene sea-level fluctuations on the evolution of tropical marine taxa. Journal of Biogeography 42: 25–38

Martin AP, Palumbi SR (1993) Body size, metabolic-rate, generation time, and the molecular clock. Proc. Natl. Acad. Sci. USA. 90: 4087–4091

Miller EF, Manica A, Amos W (2018) Global demographic history of human populations inferred from whole mitochondrial genomes. Royal Society open science 5: 180543

Normile D (2009) Bringing coral reefs back from the living dead.

Orlando L, Cooper A (2014) Using ancient DNA to understand evolutionary and ecological processes. Annual review of ecology, evolution, and systematics 45: 573–598

Paulay G (1990) Effects of late Cenozoic sea-level fluctuations on the bivalve faunas of tropical oceanic islands. Paleobiology 16: 415–434

Perry CT, Alvarez-Filip L, Graham NA, Mumby PJ, Wilson SK, Kench PS, Januchowski-Hartley F (2018) Loss of coral reef growth capacity to track future increases in sea level. Nature 558: 396–400

Pratchett MS, Caballes C, Rivera-Posada JA, and Sweatman HPA (2014) Limits to understanding and managing outbreaks of crown of thorns starfish. Oceanography and Marine Biology: An Annual Review 52: 133–200

Pratchett MS, Caballes CF, Wilmes JC, Matthews S, Mellin C, Sweatman H, Bos AR (2017) Thirty years of research on crown-of-thorns starfish (1986–2016): scientific advances and emerging opportunities. Diversity 9: 41

Rambaut A, Drummond AJ, Xie D, Baele G, Suchard MA (2018) Posterior summarization in Bayesian phylogenetics using Tracer 1.7. Systematic biology 67: 901

Ramos-Onsins SE, Rozas J (2002) Statistical properties of new neutrality tests against population growth. Molecular biology and evolution 19: 2092–2100

Shapiro B, Ho SY, Drummond AJ, Suchard MA, Pybus OG, Rambaut A (2011) A Bayesian phylogenetic method to estimate unknown sequence ages. Molecular biology and evolution 28: 879–887

Veron J, Stafford-Smith M, DeVantier L, Turak E (2015) Overview of distribution patterns of zooxanthellate Scleractinia. Frontiers in Marine Science 1: 81

Vogler C, Benzie J, Barber PH, Erdmann MV, Sheppard C, Tenggardjaja K, Wörheide G (2012) Phylogeography of the crown-of-thorns starfish in the Indian Ocean. PloS one 7: e43499

Vogler C, Benzie JAH, Tenggardjaja K, Barber PH, Wörheide G (2013) Phylogeography of the crown-of-thorns starfish: genetic structure within the Pacific species. Coral Reefs 32: 515–525

Warrick R, Oerlemans J, Beaumont P, Braithwaite RJ, Drewery DJ, Gornitz V, Lingle CS (2012) Chapter 9: Sea Level Rise. IPCC: Fifth Assessment Report (AR5)

Waelbroeck C, Labeyrie L, Michel E, Duplessy JC, McManus JF, Lambeck K, Labracherie M (2002) Sea-level and deep-water temperature changes derived from benthic foraminifera isotopic records. Quaternary Science Reviews 21: 295–305

Wu CI, Li WH (1985) Evidence for higher rates of nucleotide substitution in rodents than in man. Proc. Natl. Acad. Sci. USA. 82:1741–1745

