## Supplementary information for "Reef development and Sea level changes drive *Acanthaster* Population Expansion in the Indo-Pacific region"

**P.C. Pretorius[1], T.B. Hoareau[1,2]**

[1] Molecular Ecology and Evolution Program Marine working group, Department of Biochemistry, Genetics and Microbiology, University of Pretoria [2] Reneco International Wildlife Consultants LLC, Sky Tower, office 3902 and 3903 - Al Reem Island P.O. Box 61741 - Abu Dhabi, United Arab Emirates

### Supporting Information

**Table S1.1** FASTSIMCOAL 2 simulation parameters based on a demographic proxy that uses RSL (Waelbroeck et al. 200) as input. The new size is given relative to a modern population size given in the input of the FASTSIMCOAL file. Only changes in effective population size were altered in this case but other values must have 0 as a spacer.

| Time | | | Migrants | New size | new growth rate | migr. |
|---|---|---|---|---|---|---|
| 0 | 0 | 0 | 0 | 0.01 | 0 | 0 |
| 500 | 0 | 0 | 0 | 0.01 | 0 | 0 |
| 2000 | 0 | 0 | 0 | 0.0099724 | 0 | 0 |
| 3500 | 0 | 0 | 0 | 0.0097816 | 0 | 0 |
| 5000 | 0 | 0 | 0 | 0.0095415 | 0 | 0 |
| 6500 | 0 | 0 | 0 | 0.0091292 | 0 | 0 |
| 8000 | 0 | 0 | 0 | 0.0083342 | 0 | 0 |
| 9500 | 0 | 0 | 0 | 0.006341 | 0 | 0 |
| 11000 | 0 | 0 | 0 | 0.0043082 | 0 | 0 |
| 12500 | 0 | 0 | 0 | 0.0028395 | 0 | 0 |
| 14000 | 0 | 0 | 0 | 0.0018923 | 0 | 0 |
| 15500 | 0 | 0 | 0 | 0.0013966 | 0 | 0 |
| 17000 | 0 | 0 | 0 | 0.0011802 | 0 | 0 |
| 18500 | 0 | 0 | 0 | 0.0010568 | 0 | 0 |
| 20000 | 0 | 0 | 0 | 0.001 | 0 | 0 |
| 21500 | 0 | 0 | 0 | 0.0010481 | 0 | 0 |
| 23000 | 0 | 0 | 0 | 0.0011045 | 0 | 0 |
| 24500 | 0 | 0 | 0 | 0.0012065 | 0 | 0 |
| 26000 | 0 | 0 | 0 | 0.0013699 | 0 | 0 |
| 27500 | 0 | 0 | 0 | 0.0015814 | 0 | 0 |
| 29000 | 0 | 0 | 0 | 0.0019239 | 0 | 0 |
| 30500 | 0 | 0 | 0 | 0.0021664 | 0 | 0 |
| 32000 | 0 | 0 | 0 | 0.002302 | 0 | 0 |
| 33500 | 0 | 0 | 0 | 0.0022705 | 0 | 0 |
| 35000 | 0 | 0 | 0 | 0.0022086 | 0 | 0 |
| 36500 | 0 | 0 | 0 | 0.0023212 | 0 | 0 |
| 38000 | 0 | 0 | 0 | 0.0028676 | 0 | 0 |
| 39500 | 0 | 0 | 0 | 0.0031269 | 0 | 0 |
| 41000 | 0 | 0 | 0 | 0.0028097 | 0 | 0 |
| 42500 | 0 | 0 | 0 | 0.0025243 | 0 | 0 |
| 44000 | 0 | 0 | 0 | 0.0024292 | 0 | 0 |
| 45500 | 0 | 0 | 0 | 0.0025272 | 0 | 0 |
| 47000 | 0 | 0 | 0 | 0.0025395 | 0 | 0 |
| 48500 | 0 | 0 | 0 | 0.002503 | 0 | 0 |
| 50000 | 0 | 0 | 0 | 0.0026172 | 0 | 0 |
| 51500 | 0 | 0 | 0 | 0.0027553 | 0 | 0 |
| 53000 | 0 | 0 | 0 | 0.0034049 | 0 | 0 |
| 54500 | 0 | 0 | 0 | 0.0037722 | 0 | 0 |
| 56000 | 0 | 0 | 0 | 0.003512 | 0 | 0 |
| 57500 | 0 | 0 | 0 | 0.0032203 | 0 | 0 |
| 59000 | 0 | 0 | 0 | 0.0034925 | 0 | 0 |
| 60500 | 0 | 0 | 0 | 0.0040706 | 0 | 0 |
| 62000 | 0 | 0 | 0 | 0.0035895 | 0 | 0 |
| 63500 | 0 | 0 | 0 | 0.0024759 | 0 | 0 |
| 65000 | 0 | 0 | 0 | 0.0020505 | 0 | 0 |
| 66500 | 0 | 0 | 0 | 0.0020616 | 0 | 0 |
| 68000 | 0 | 0 | 0 | 0.0023201 | 0 | 0 |
| 69500 | 0 | 0 | 0 | 0.002394 | 0 | 0 |
| 71000 | 0 | 0 | 0 | 0.0025613 | 0 | 0 |
| 72500 | 0 | 0 | 0 | 0.0028368 | 0 | 0 |
| 74000 | 0 | 0 | 0 | 0.0043575 | 0 | 0 |

| | | | | | | |
|---|---|---|---|---|---|---|
| 75500 | 0 | 0 | 0 | 0.00458409 | 0 | 0 |

**Table S1.2** Nucleotide accession numbers of individuals used in each of the five populations of COTS throughout the Indo-Pacific region. This includes *CR* and *COI* depending on availability

| Population/Marker | **EMBL Accession**<br>CR | |
|---|---|---|
| *A. solarius* (Western Pacific) | HE612183-HE612221,<br>HE612339-HE612303,<br>HE612318-HE612649,<br>HE612660-HE612670,<br>HE612706-HE612768,<br>HE612772-HE612822 | |
| *A. solarius* (Central Pacific) | HE612651-HE612656,<br>HE612671-HE612704,<br>HE612823-HE612855 | |
| Population/Marker | CR | COI |
| *A. planci* (North Western Indian Ocean) | HE608415-HE608439 | FM174564,<br>FM174574-FM174592 |
| *A. planci* (North Eastern Indian Ocean) | HE608415-HE608509 | FM174545-FM174592 |
| *A. mauritiensis* | HE608326-HE608370,<br>HE608375-HE608408 | FM174597-FM174637 |

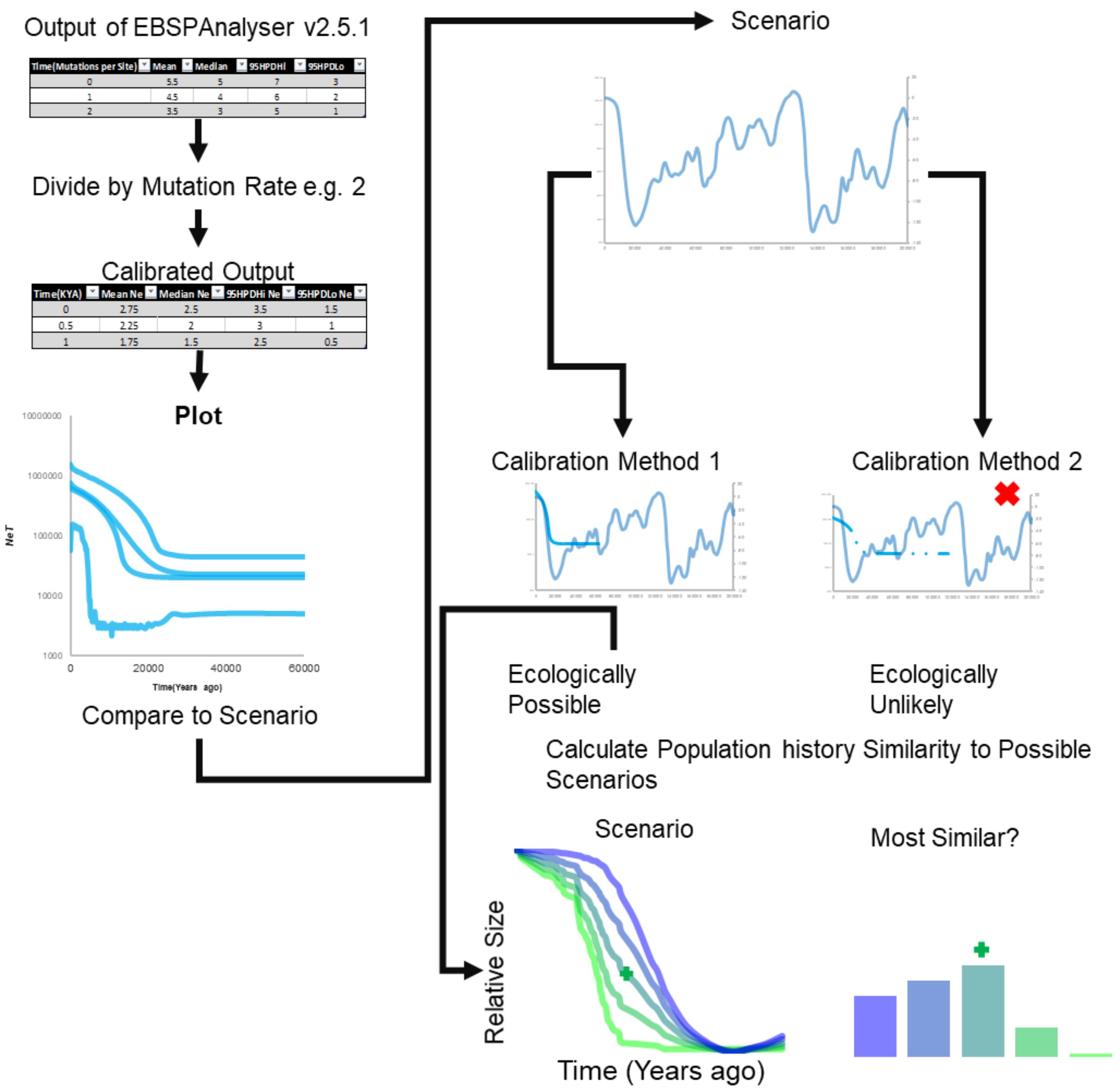


**Fig. S1.1** Conceptual approach of the new calibration method and its validation. This includes generating EBSP plots in BEAST2, followed by conversion of mutations per site to years before present. This output is generated via the use of multiple calibration methods. The ecologically most likely scenario is selected based on expansions before or after the LGM (19-26 KYA). The EBSP is then compared to a gradient of scenarios between the most likely factors to influence reef communities. The highest similarity indicates the most likely scenario.

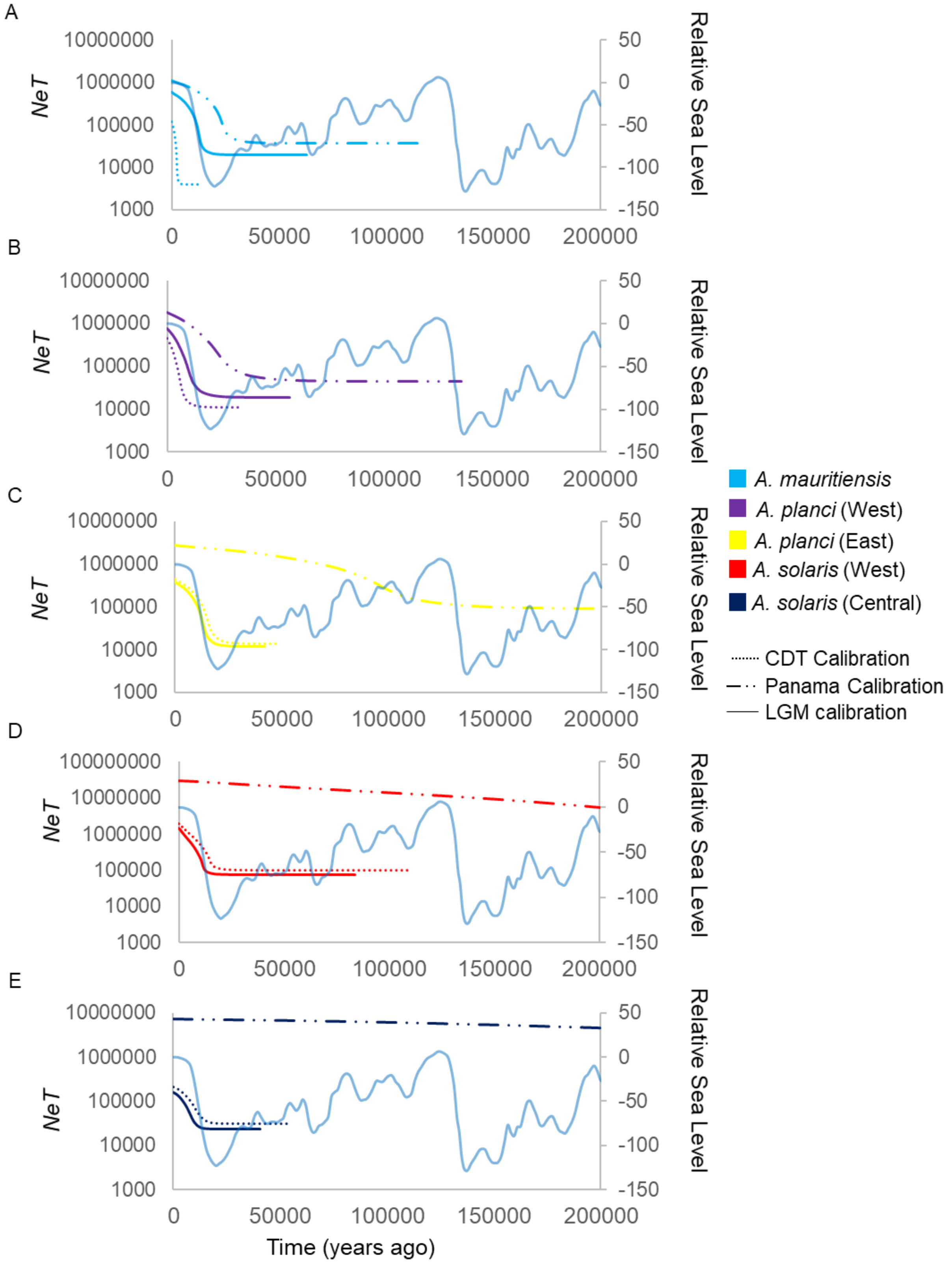


**Fig. S1.2** Median values of extended Bayesian skyline plots for the five selected COTS populations. Darkest lines indicate LGM calibrated histories. Lightest lines indicate inferences calibrated using substitution rates derived from the biogeographic calibration (Lessios 2008). The intermediate shading in the Pacific Ocean populations indicate inferences calibrated using mutation rates obtained using the CDT calibrated demographic history.

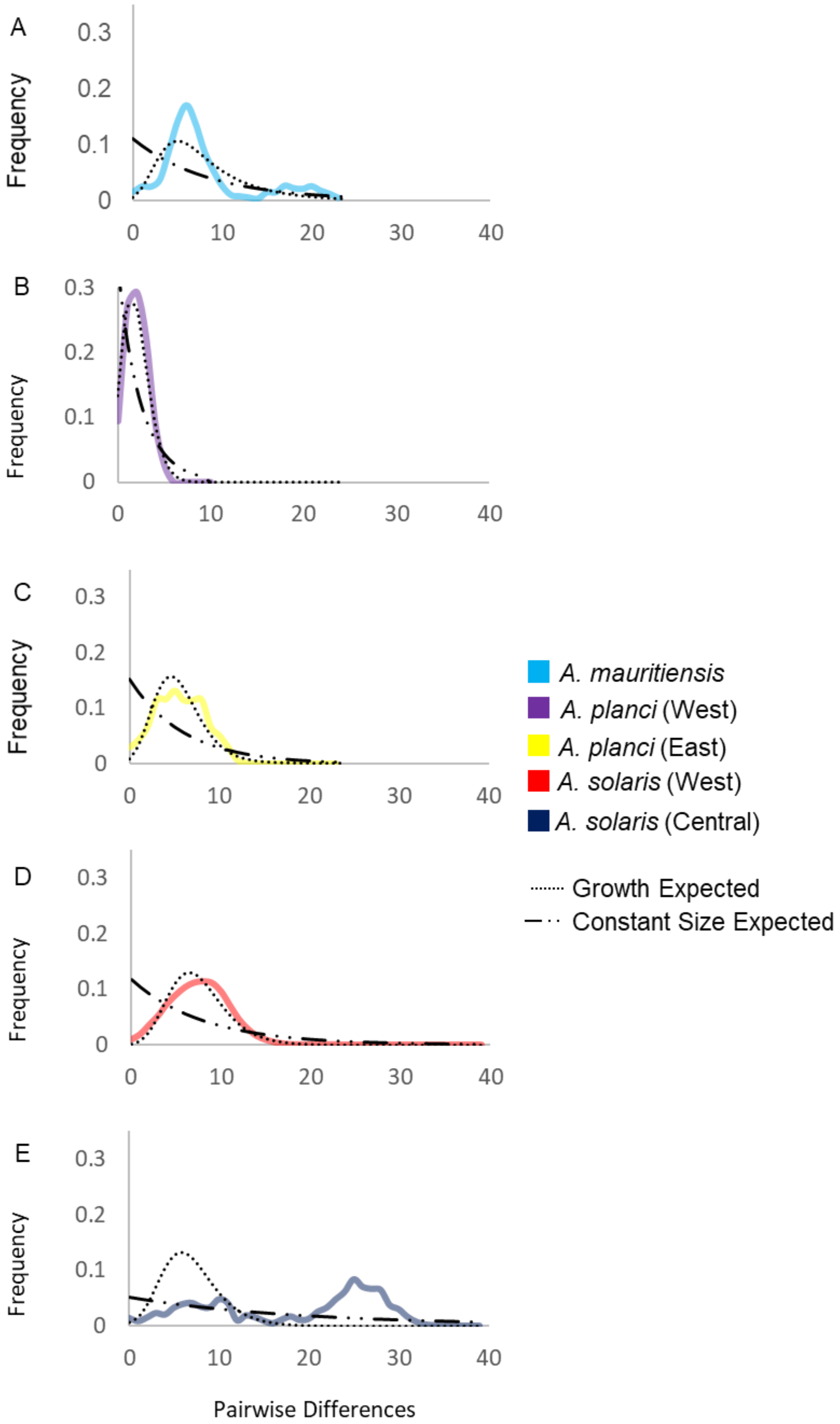
A
B
C
D
E
Frequency
0.3
0.2
0.1
0
0
10
20
30
40
Pairwise Differences
A. mauritiensis
A. planci (West)
A. planci (East)
A. solaris (West)
A. solaris (Central)
Growth Expected
Constant Size Expected

**Fig. S1.3** Mismatch distribution plots of the 5 identified populations of COTS. The default scenario of constant population size is indicated by a dashed black line. The green line represents the smoothed empirical data. The histogram plots indicate the actual counts of distance categories of the empirical data

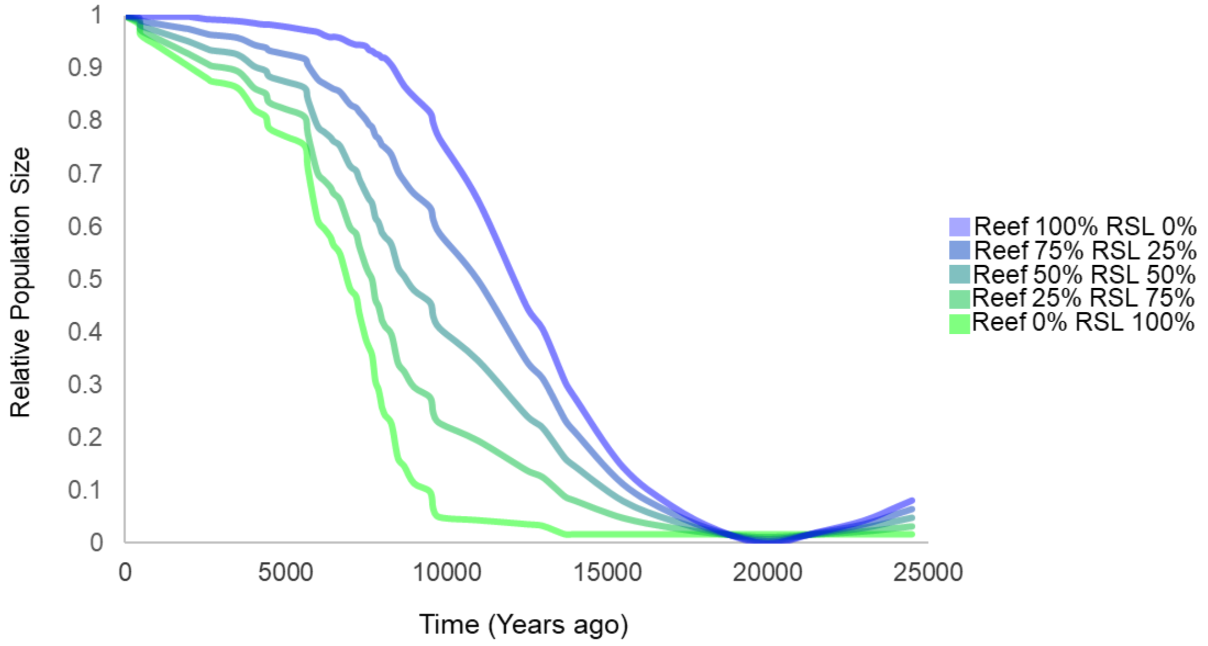


**Fig. S1.4** Gradient of possible scenarios with changing relative contributions from reef development and sea-level rise to increases in population size of populations linked to the present study. These factors are assumed to be the most important contributory factor to reef communities in this study. The scenarios will serve as the standard to compare calibrated Bayesian skyline plots to identify the scenario with the highest similarity to the demographic histories.
